# Shade enhancers, a novel class of redundant shadow enhancers uncovered by functional JAK-STAT signalling pathway dissection in *Drosophila*

**DOI:** 10.64898/2026.09.28.754901

**Authors:** C. Sánchez-Higueras, L. Terrón-González, D. Barcenilla-Merino, M.S. García-Ferrés, Á. Marrufo-Mena, S. Sotillos, J. C.-G Hombría

## Abstract

JAK-STAT signalling pathway activation relies mostly on the transcriptional regulation of its ligands. The pathway, conserved in vertebrates and invertebrates, is required for the development and homeostasis of various organs. In contrast to the complexity of the vertebrate pathway, with a multiplicity of ligands, receptors, kinases and transcription factors, the simplicity of the *Drosophila* pathway with only three ligands (Upd1, Upd2 and Upd3), a single receptor (Dome), a JAK kinase (Hop) and a STAT transcription factor (STAT92E) facilitates the genetic study of its multiple functions. To discover how, when and where the Upd1 ligand performs each organogenetic function, we have searched for the different enhancers controlling its dynamic spatio-temporal regulation and used them to generate minigenes replicating specific stages of Upd1 expression. The combination of these minigenes with mutations deleting all three Upd ligands, allowed us to identify the timing when the various embryonic functions are mediated, including head involution, tracheae, *corpora allata* and prothoracic gland specification, posterior spiracle formation and hindgut elongation. We uncover most of the enhancers controlling STAT activation during embryonic organogenesis and those driving activation in the adult testis and the ovary. As expected, we identified redundant shadow enhancers but, interestingly, we found a different type of redundant enhancers that mediate identical developmental functions although driving completely different expression patterns. We name this new class of functionally redundant enhancers driving different expression as *shade enhancers*. We propose shade enhancers provide evolutionary plasticity as they could allow the pattern of gene expression to diversify without altering the underlying body plan.

## Introduction

JAK-STAT signalling is a conserved pleiotropic pathway required in vertebrates and invertebrates for many different developmental and homeostatic functions. Although the basic signalling pathway elements are conserved, the reduced number of gene duplications makes the invertebrate signalling pathway more streamlined, offering an ideal system to genetically analyse JAK-STAT functions, especially those involved in development and morphogenesis (Hombria and Sotillos, 2013). The three *Drosophila* ligands genes, *unpaired1* to *unpaired3* (*upd1-3*), are clustered in the X chromosome (Fig.1A). Upd binding, activates homodimers of a single receptor (Domeless) that through a single JAK kinase (Hopscotch) phosphorylate the only STAT transcription factor (STAT92E) [Reviewed in (Hombria and Brown, 2002)]. Embryos homozygous for the *Deficiency os1A* (*Dfos1A*) deleting simultaneously the *upd1*, *upd2* and *upd3* ligands have defects in segmentation, cephalisation, hindgut, foregut, posterior spiracles, trachea, endocrine glands and tentorium development, which are indistinguishable from the phenotypes caused by the complete abrogation of the maternal and zygotic products of the receptor, the JAK kinase or the STAT transcription factor of the pathway (Binari and Perrimon, 1994; Brown et al., 2001; Harrison et al., 1998; Hou et al., 1996; Yan et al., 1996). Previous functional analyses indicated that *upd1* and to a minor extent *upd2* are required for embryo development, while *upd2* and *upd3* are mostly involved in larval and adult homeostasis (Hombria et al., 2005). Accordingly, *upd2, upd3* and the double *upd2 upd3* mutants give rise to viable and fertile *Drosophila* adults (Osman et al., 2012; Wang et al., 2014), while *upd1* mutants are embryonic lethal (Harrison et al., 1998). These results indicate that Upd1 can mediate most, if not all developmental functions, as confirmed by the strong and defined embryonic expression of *upd1* (Fig.1B).

**Fig. 1.**
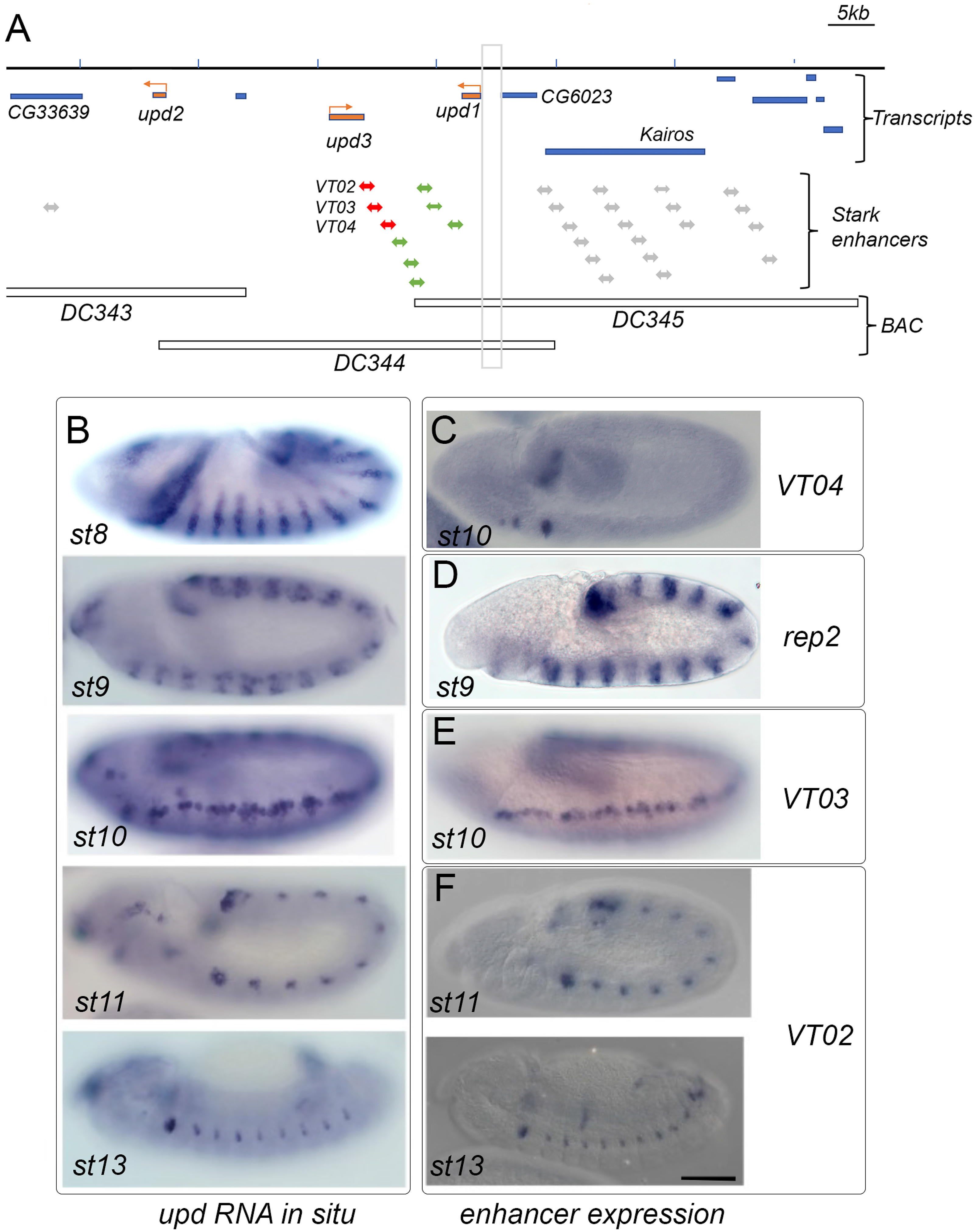
*upd* locus and expression of *upd* RNA and *cis*-regulatory elements. (A) Scheme showing *upd1-3* transcription units as orange arrows. Stark lab *cis*-regulatory regions shown as double arrows with those not in the locus highlighted in grey, in red the ones analysed in this study, and in green those that we did not. Grey vertical box highlights the upstream region analysed to make *rep1* and *rep2* reporters. Horizontal boxes show the extent of BAC duplications used in this study. (B) *D. melanogaster upd* RNA expression at the indicated stages. (C-F) RNA *in situ* of mCherry or GFP in various reporter genes. (C) *VT04-mCherry* expression in tentorium/gland primordia and hindgut. (D) *rep2-GFP* expression in segmental stripes. (E) *VT03-mCherry* expression along the lateral epidermis prior to tracheal specification. (F) *VT02-mCherry* expression in tracheal cells at two stages of development. Scale bar 50 µm.

The dynamic spatio-temporal pattern of *upd* transcription prevents knowing at what stage STAT92E activation controls each of its developmental functions. Thus, isolating the specific enhancers regulating the expression of *upd* can provide tools to dissect specifically the timing of the diverse pathway’s functions. Such tools could be useful for manipulating specific functions during embryonic organogenesis, stem cell regulation in blood, gut, ovary and testis during larval and adult stages, as well as understanding its physiological functions during regeneration and natural immunity (Amoyel and Bach, 2012; Herrera and Bach, 2019; Hombria and Sotillos, 2013; Yu et al., 2022).

To discover the temporal windows when the JAK-STAT pathway is required for its diverse developmental functions, we isolated specific enhancers controlling *upd1* embryonic expression and generated *upd1* cDNA minigenes. Combining each minigene with a deletion of all three Upd ligands, we were able to rescue the formation of particular organs, identifying at which stage the ligand mediates the different pathway’s functions during organogenesis. Our experiments uncover the existence of a new class of redundant enhancers we named *shade enhancers*, that differ from the previously described shadow enhancers in driving completely different patterns of expression. We propose *shade enhancers*, which can only be discovered using functional analyses, probably exist in many other genes providing developmental plasticity that may have important evolutionary implications.

## Results

The *upd* locus is located in a 90kb region on the X chromosome which is completely deleted by the deficiency *Dfos1A* (Hombria et al., 2005). To broadly define the enhancers’ locations, we tested the capacity of three overlapping Bacterial Artificial Chromosomes (BAC) to rescue *Dfos1A* embryos (Fig. 1A). The DC343 BAC, containing the *upd2* transcription unit and approximately 10kb of its upstream region, was unable to rescue any of the cuticular embryonic defects, suggesting that although Upd2 can weakly compensate *upd1* mutant phenotypes (Hombria et al., 2005), the absence of embryonic *cis*-regulatory elements prevents its activation. Both DC344, containing the *upd3* and *upd1* transcription units, and DC345, containing the *upd1* transcription unit, were able to rescue all *Dfos1A* embryonic phenotypes. However, these animals did not reach adulthood, suggesting the DC344 and DC345 lack essential *upd* larval and pupal functions (see Materials and methods), although we cannot completely discard the lethality to be caused by associated mutations in the X chromosome.

### Identification of organogenetic *upd* regulatory elements

To locate the embryonic regulatory elements in this large locus, we took advantage of the information available in the <u>Stark lab</u> database. This public database reports the embryonic expression of Gal4 driven by thousands of overlapping 2kb *Drosophila* genome segments (Kvon et al., 2014) including many located in the *upd3*-*upd1* intergenic region (Fig.1A, double-head arrows). This allowed us to locate several enhancers active during the organogenesis period (6-16 hours of development), which replicate most aspects of the *upd1* RNA expression. To determine the precise temporal activity of these enhancers, we made *mCherry* and *GFP* reporters and analysed their expression (Fig.1C-F). We studied the enhancer activity using RNA *in situ* rather than analysing the reporter protein expression to avoid the confounding effect of the GFP and mCherry protein stability. We found that from stage 9 (st9) to stage 12 (st12) the VT04 enhancer (VT637<u>04</u>) drives expression in the hindgut, and from st10 to st12 it drives expression in the intercalary, maxillary and labial cephalic segments in the primordia of the future endocrine glands and the tentorium [Fig. 1C, and (Barcenilla-Merino et al., 2025)]. VT03 (VT637<u>03</u>) drives very transient expression at st10 in an almost continuous antero-posterior stripe of cells extending from the mandibular to the last abdominal segment (Fig.1E). After st12 most VT03 activity stops, remaining exclusively in the anterior spiracle primordium. Considering their morphology, size and position at later stages when analysing the more stable GFP and mCherry proteins, this reporter is expressed in the primordia of the most dorsal neuroblasts and in the cells that will give rise to the tracheal ventral branches (Sup. Fig.1A-C) (Manning and Krasnow, 1993; Schmidt et al., 1997). VT02 (VT637<u>02</u>) drives expression in a subset of tracheal pit cells that will form the spiracular branch (st11-14), in the anterior (st11-17) and in the posterior spiracle primordia (st11-13), in the hindgut (st11-13), the midgut (st15), and from st12 in the tentorium (Fig.1F and Sup. Fig. 1E-G). We subdivided VT02 in two overlapping elements (VT02A, VT02B), both of which gave the same expression in the trachea and spiracles suggesting that these enhancers locate to the overlapping 200bp region (Sup. Fig. 2). Subdivision of VT03 locates the antero-posterior stripe enhancer to VT03B, a 740bp region, and dissection of VT04 mapped head and hindgut enhancers to VT04.2 a 793bp region. We did not analyse in detail other VT elements reported in the Stark database, one of which shows *upd1*-like expression in the proventriculus, the hindgut and anterior spiracles (VT63708), and another expressed in the procephalic ectoderm (VT63713).

We also made reporter constructs of *upd1* with the upstream *cis* regulatory region that is not covered by the Stark’s genome analysis. As the overall *upd1* RNA embryonic transcription is conserved in *D. virilis* and *D. pseudobscura* (Sup. Fig. 3), we used the upstream sequence conservation as a guide to generate two reporter constructs, *rep1-GFP* and *rep2-GFP* (Sup. Fig. 2). While *rep1* does not drive embryonic expression, *rep2* reproduces *upd1*’s st9 to st10 segmental expression (Fig. 1D).

To test if all the above reporters contain enhancers active at later development, we dissected larva 3 (L3) imaginal discs, adult testis and ovary. We did not detect expression in the imaginal discs with any of these reporters. However, all five constructs are active at the testis terminal region in the mesodermal Head Cyst Cells (HCC) holding the sperm heads (Desai et al., 2009), indicating the presence of at least four HCC redundant shadow enhancers (Fig. 2A-F). VT03 also drives expression in the testis hub, which constitutes the germ cell niche, and in the mesodermal Cyst Cells (CC) surrounding early-stage spermatocytes (Fig. 2G) (Fuller, 1993). In the ovary (Fig. 2H), VT03 drives expression in the germarium restricted to the mesodermal follicle cells, and in the vitellarium it labels both the migrating polar and border cells (Spradling, 1993). The location of the gonadal regulatory elements can be narrowed down to a shorter 609bp VT03A fragment (Sup. Fig. 2).

**Fig. 2.**
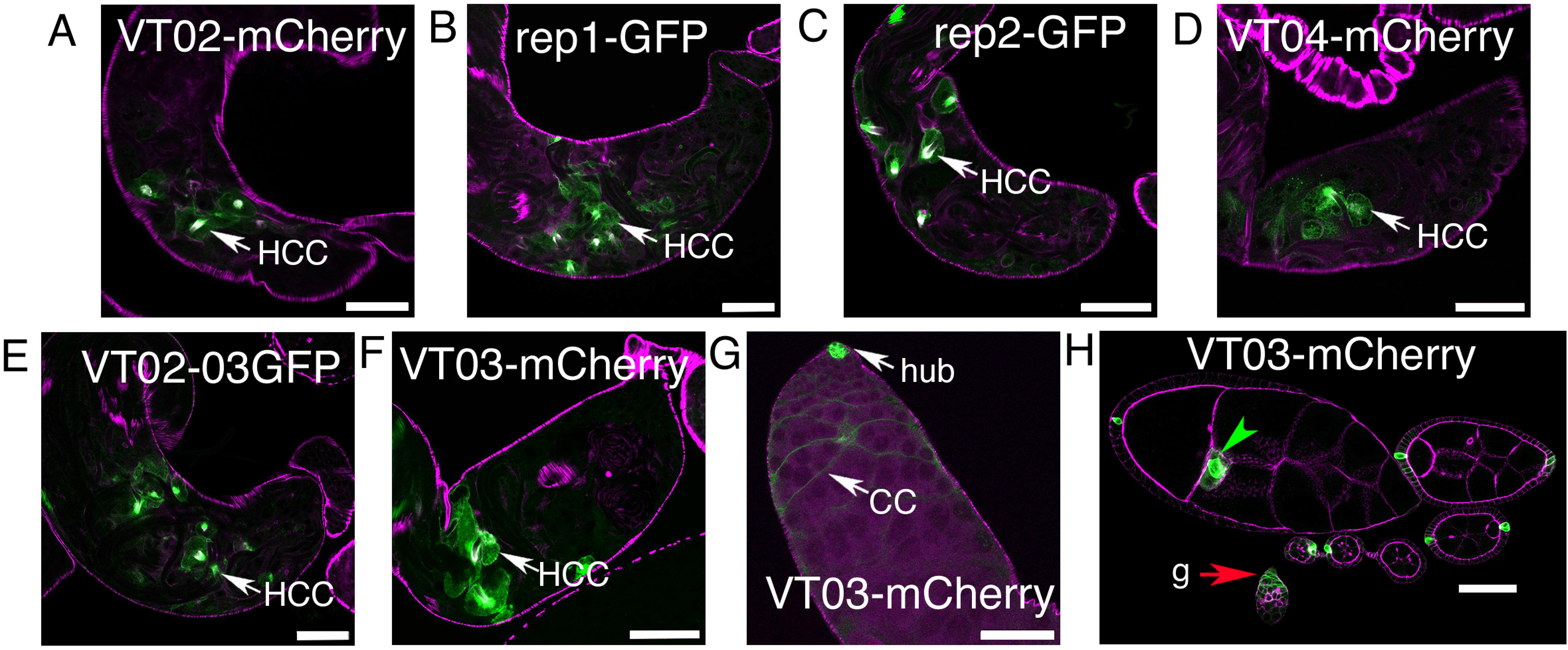
Testis and ovary expression of *upd cis*-regulatory elements. Antibody stainings detecting reporter expression in gonads. (A-F) Testis terminal region showing expression in the Head Cyst Cells (HCC, white arrows) driven by all tested reporters. (G-H) *VT03-mCherry* expression in the gonads. In the testis anterior region (G) *VT03-mCherry* is active in the hub and in the cyst cells (CC). In the ovary (H) *VT03-mCherry* is active in the germarium (g, red arrow) where it labels the follicle mesodermal cells protecting the germ cells. In the vitellarium, high levels of expression are observed in the polar cells and lower levels in the border cells surrounding them (green arrowhead). Expression in the border cells is probably due to mCherry perdurance from the earlier expression in the germarium. Scale bars 50 µm.

Although our results have located enhancers recapitulating all sites of *upd1* expression in the adult gonads and most enhancers required for *upd1* expression during embryonic organogenesis, other regions not identified in this study must contain additional shadow enhancers, as indicated by the fact that DC345 which lacks VT02, VT03 and VT04 (Fig. 1A) can rescue all embryonic morphogenetic functions.

### Generation of *upd* minigene rescue constructs

To discover what are the organogenetic functions that Upd1 signalling mediates through the *VT02*, *VT03*, *VT04* and *rep2* enhancers, we made constructs where *upd1* cDNA fused C-terminally to an HA-tag is under the regulation of each of these elements (Materials and methods). The HA tag in these constructs is inserted downstream of putative furin sites that, if functional, would cleave the tag before Upd is secreted (Takemura et al., 2021). We were able to detect strong HA signal in VT02 and VT04 minigenes (Sup. Fig. 4), but not in the VT03 or rep2 minigenes, probably because of their brief expression and the rapid degradation of the intracellular cleaved C-terminal element.

To analyse if the minigenes are functional, we tested their ability to rescue *Dfos1A* embryo cuticle defects. To that end we first checked if VT02, VT03, VT04 and rep2 enhancer expression is maintained in *Dfos1A* embryos (Sup. Fig. 5). *Dfos1A* embryos are lethal, showing penetrant defects where some segments are fused or deleted due to JAK-STAT requirement for pair-rule gene expression (Hou et al., 1996; Small et al., 1996; Yan et al., 1996). This results in a characteristic cuticle phenotype where many segments are frequently deleted or fused with the exception of A1, A2 and A3 that tend to be present (Fig. 3A-B). Besides this, in *Dfos1A*embryos the cephalopharyngeal skeleton is abnormal, the tracheae do not form and the posterior spiracles are stunted (Fig. 3B’-B’’). Despite these abnormalities we observe that all four enhancers are active in *Dfos1A* embryos (Sup. Fig. 5), indicating that the minigenes can be used to rescue the associated defects.

**Fig. 3.**
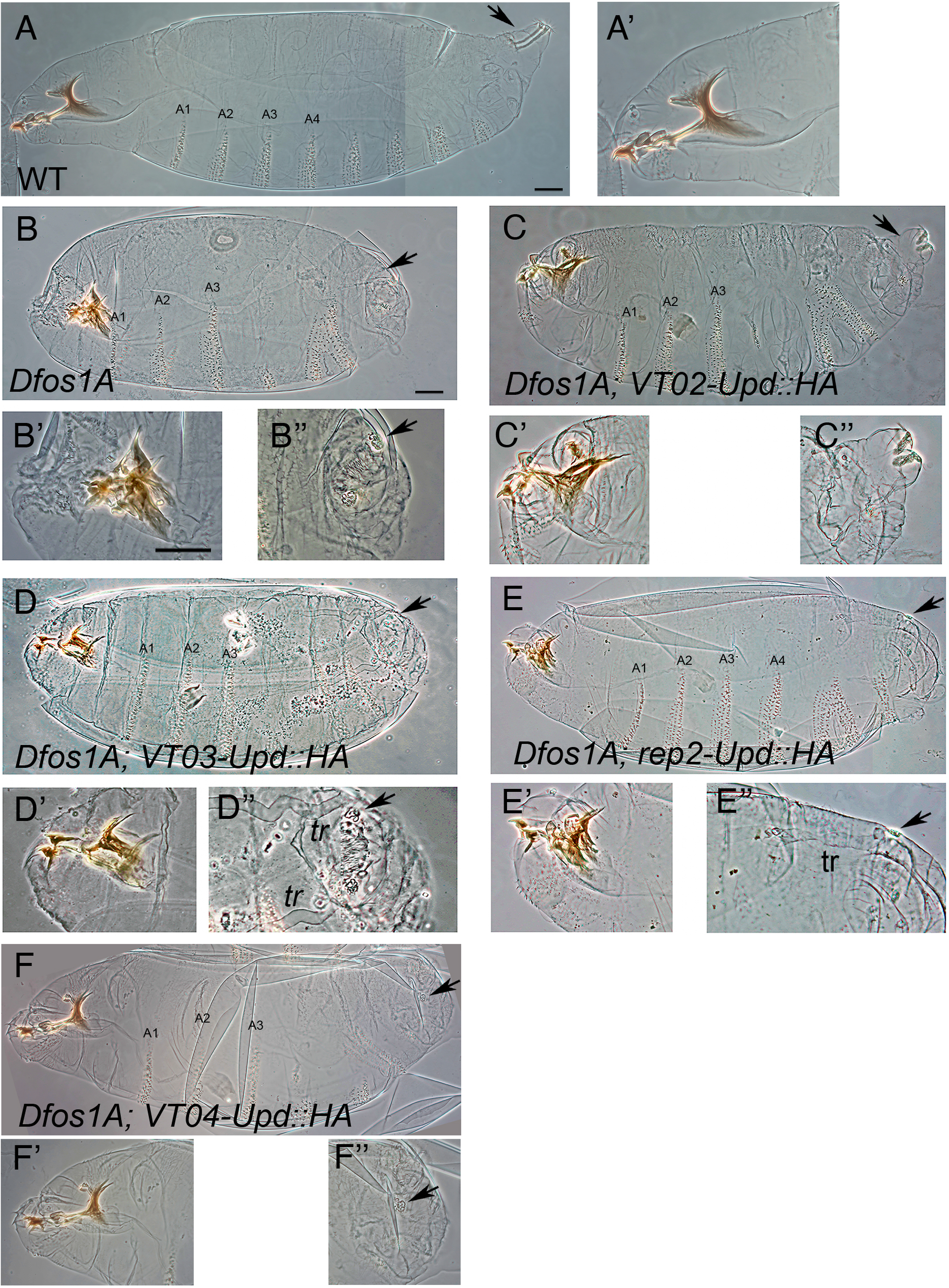
Minigene cuticular rescues in *upd* null embryos. (A-F) Phase contrast images of late embryo cuticles with close ups of the cephalic (’) and tail regions (’’) to highlight the cephalopharyngeal skeleton and the posterior spiracle formation. (A) Wild type embryo. (B) *Dfos1A* embryo showing segmental defects, lack of trachea, abnormal mouth parts (B’), and stunted posterior spiracles (B’’). (C) *Dfos1A VT02-Upd::HA* embryos develop a filzkörper filter and a protruding stigmatophore in the posterior spiracle (C’’). (D) *Dfos1A VT03-Upd::HA* embryos form the dorsal tracheal trunk (tr) but do not rescue posterior spiracle structures. (E) *Dfos1A rep2-Upd::HA* embryos also form the dorsal tracheal trunk but do not rescue posterior spiracle structures. (F) *Dfos1A VT04-Upd::HA* embryos develop an almost normal cephalopharyngeal skeleton. The position of abdominal segments 1 to 4 (A1-A4) are labelled when present. Black arrows point at the site where the posterior spiracles develop. Scale bar 50 µm.

With the exception of *rep2-Upd::HA* that significantly recovered the formation of the A4 segment (Fig. 3E), none of the other minigenes rescued *Dfos1A* segmentation defects, as expected from their expression being active from st10 onwards when the segmentation process is complete. However, we found that each minigene rescued particular aspects of the *Dfos1A* defects (Table 1). In *Dfos1A VT02-Upd::HA* the posterior spiracles recover the internal filter (filzkörper) and form the characteristic protruding structure of the stigmatophore, while other aspects like the head defects or the trachea formation are not rescued (Fig. 3C). In contrast, *Dfos1A VT03-Upd::HA* cuticles form a trachea, but do not rescue the posterior spiracle structure (Fig. 3D). Similarly, *Dfos1A rep2-Upd::HA* embryos also rescue trachea formation, but not head nor posterior spiracles (Fig. 3E). Finally, *Dfos1A VT04-Upd::HA* embryos present a normalized cephalopharyngeal skeleton but do not rescue the formation of the trachea nor the posterior spiracles (Fig. 3F). These results demonstrate that the four minigene constructs are functional and able to rescue particular aspects of JAK-STAT signalling.

**Table 1:** Summary of temporal enhancer expression and minigene embryonic rescues.

| Enhancer | Stages | Embryonic expression | Testis | Ovary | Minigene rescue |
| --- | --- | --- | --- | --- | --- |
| VT02 | 11-17 | spiracular branch (st11-14),<br>anterior spiracular primordia (st11-17),<br>posterior spiracular primordia (st11-13),<br>hindgut (st11-13),<br>midgut (st15)<br>tentorium (st12-17) | HCCs | no expression | Posterior spiracles |
| VT03 | 10-12 | dorsal neuroblasts (st10)<br>ventral tracheal branch bud (st10)<br>anterior spiracle primordium (st12) | HCCs<br>and<br>hub | Mesodermal cells<br>and germarium | Endocrine glands<br>Tracheae |
| VT04 | 9-12 | hindgut (st9-12)<br>endocrine glands and tentorium (st10-12) | HCCs | no expression | Head formation<br>Endocrine glands<br>Hindgut |
| rep1 | - | no expression | HCCs | no expression | - |
| rep2 | 9-10 | weak blastoderm (st5-6)<br>segmental expression (st9-10)<br>hindgut (st8-10) | HCCs | no expression | Tracheae<br>Hindgut elongation |

### Rescue of internal organs by different minigene constructs

To study in more detail the formation of internal organs in the rescue experiments, we stained the embryos with specific reporters or antibodies labelling the trachea, the endocrine glands (*corpora allata* and prothoracic glands) and the posterior spiracles, all of which do not form in *Dfos1A* embryos (Brown et al., 2001; Garcia-Ferres et al., 2022; Sanchez-Higueras et al., 2014). Consistent with the cuticle observations, double staining with anti-Sal and anti-En antibodies shows that in *Dfos1A, VT02-Upd::HA* embryos, the posterior spiracles develop forming their characteristic circle of En expression around the posterior spiracle opening which is composed of anterior and posterior compartment cells, while the tracheal dorsal trunk does not form (Fig. 4A-B and Table 1). This contrasts with *Dfos1A, VT03-Upd::HA* embryos where Engrailed spiracle expression is absent from the anterior A8 compartment, while the tracheal dorsal trunk is formed (Fig. 4C), and a similar rescue phenotype is observed in *Dfos1A, rep2-Upd::HA* embryos (Fig. 4D).

**Fig. 4.**
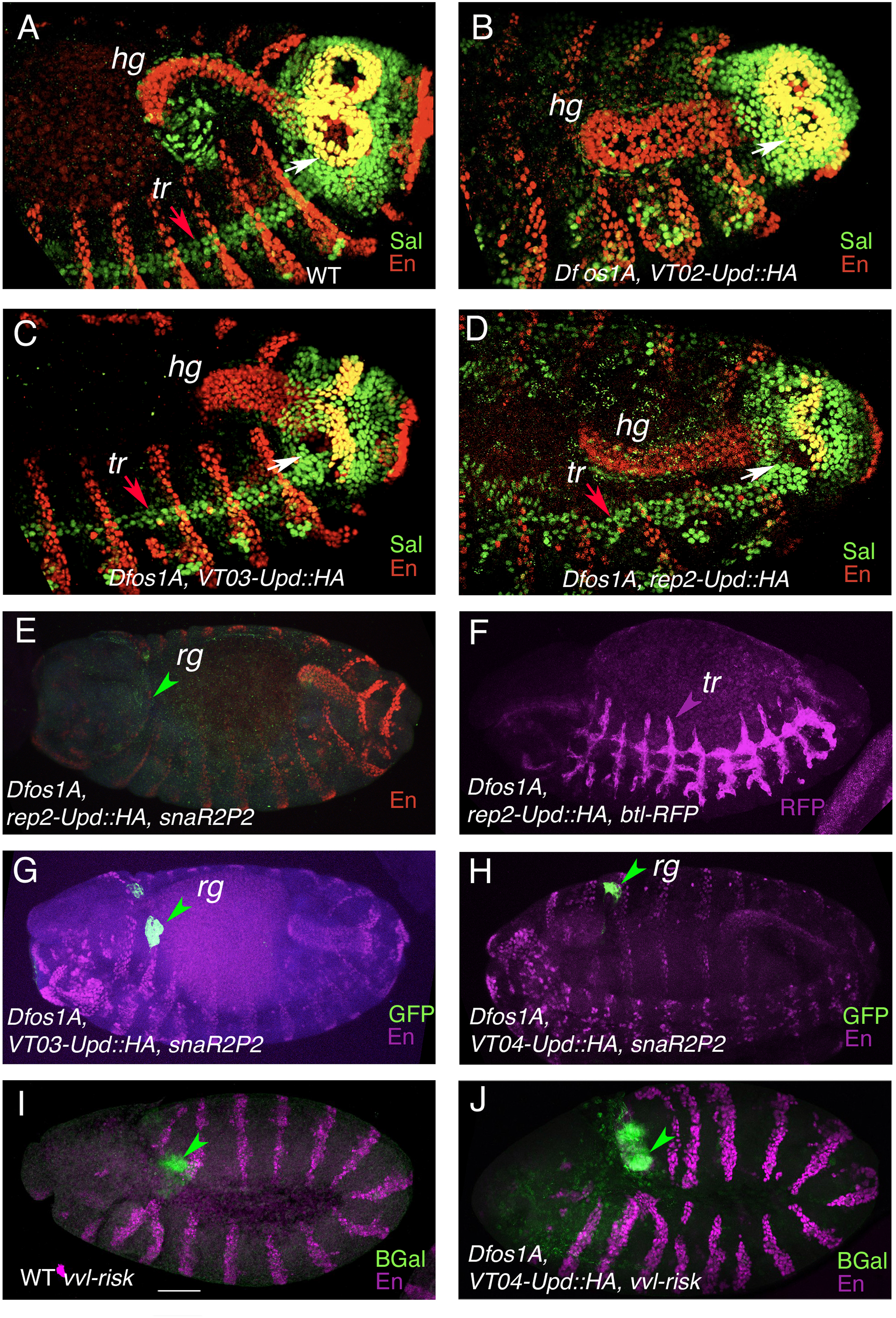
Minigene internal organ rescues of *upd* null embryos. (A-D) Posterior abdomens of stage 14 embryos stained with anti-En (red) and anti-Sal (green). In wild type embryos (A) Engrailed and Spalt are co-expressed in the posterior spiracles forming a circle in the cells that will form the protruding external part (the stigmatophore, co-expression is detected as a yellow circle, white arrow). Sal also labels the tracheal dorsal trunk (tr, red arrow) and Engrailed is also expressed in the hindgut (hg) and in the posterior compartment of each segment. In *Dfos1A VT02-Upd::HA* embryos (B) the En stigmatophore circular expression is present, but the dorsal tracheal trunk is not formed, and the hindgut does not elongate. In *Dfos1A VT03-Upd1::HA* embryos (C) the formation of the dorsal tracheal trunk is rescued, but not the circular En expression in the stigmatophore nor the hindgut extension. In *Dfos1A rep2-Upd1::HA* embryos (D) the formation of the dorsal tracheal trunk is rescued, as well as hindgut elongation but not the circular En expression in the stigmatophore. *Dfos1A rep2-Upd1::HA* embryos carrying the *snaR2P2-GFP* gland reporter gene (E) show lack of *corpora allata* and prothoracic gland primordia (rg, green arrowhead). *Dfos1A rep2-Upd1::HA* embryos carrying the *btl-moeRFP* tracheal reporter gene (F) show an almost complete rescue of the trachea. *Dfos1A VT03-Upd1::HA* embryos carrying the *snaR2P2-GFP* reporter gene (G) show a rescue of the *corpora allata* and prothoracic gland primordia specification, and a similar rescue is observed in *Dfos1A VT04-Upd1::HA snaR2P2-GFP* embryos (H). *Dfos1A VT04-Upd1::HA* embryos also have a rescued hindgut as shown by the elongated En expression. Similar expression of the *vvl-risk-lacZ* hindgut reporter gene in wild type embryos (I) and in *Dfos1A VT04-Upd1::HA* embryos (J). Except F, all embryos are stained with anti-En. A-H embryos are at the retracted germ band stage. I-J are at the extended germ band stage. Scale bar 50 µm.

To study the formation of the *corpora allata* and prothoracic glands, we combined the rescue embryos with the gland specific *snaR2P2-GFP* reporter (Sanchez-Higueras et al., 2014). We observe that in *Dfos1A rep2-Upd::HA* embryos the minigene is unable to activate *snaR2P2-GFP* gland expression, although it can rescue almost completely trachea development (Fig. 4E-F). In contrast both *VT03-Upd::HA and VT04-Upd::HA* can rescue endocrine gland specification and migration as detected with *snaR2P2-GFP* (Fig. 4G-H).

JAK-STAT signalling is required for cell rearrangement in the hindgut (Johansen et al., 2003). In *Dfos1A VT02-Upd::HA* and *Dfos1A VT03-Upd::HA* the hindgut is shorter and wider, indicating a lack of cell rearrangement (Fig. 4B-C). This phenotype is completely rescued in *Dfos1A, rep2-Upd::HA* embryos which have early hindgut expression and a similar hindgut rescue is obtained in *Dfos1A VT04-Upd::HA* (Fig. 4D,H). To analyse at the gene expression level the hindgut rescue we studied the activation of the *vvl-risk-lacZ* reporter made with a *ventral veinless* enhancer whose activation in the hindgut ectoderm depends on JAK-STAT activity (Sotillos et al., 2010) observing that *VT04-Upd::HA* completely rescues *vvl risk-lacZ* expression in *Dfos1A* embryos (Fig. 4I-J).

### Regulation of the *upd* enhancer elements in the embryonic trachea

The JAK-STAT pathway is required for the specification of the tracheae, the tentorium and the ectodermal endocrine glands (*corpora allata* and prothoracic glands). These four organs arise from metamerically homologous cells formed in the cephalic and the trunk segments whose regional differentiation is mediated by Hox protein function, where Dfd specifies *corpora allata*, Scr prothoracic glands, and Antp, Ubx, Abd-A and Abd-B specify trachea development (Barcenilla-Merino et al., 2025; Sanchez-Higueras et al., 2014).

To start unravelling the upstream elements controlling Upd1 regulation in these organs we analysed if any of the studied *upd* organogenetic enhancers is under Hox regulation. As *VT03*-*Upd1::HA* and *VT04-Upd1::HA* minigenes are capable of rescuing gland formation, we analysed if their expression is affected in *Dfd Scr* mutants where there is no Hox functional input in the embryonic maxilla and labium and the glands are not formed. While in *Dfd Scr Abd-B VT04-mCherry* cephalic expression is absent from the head (Barcenilla-Merino et al., 2025), in *Dfd Scr Abd-B* or in *lab Dfd Scr* embryos, VT03 reporter expression is unaffected (Fig. 5B and Sup. Fig. 1C-D,H). It has been shown that in *Dfd Scr* embryos the *sna* ring gland specific enhancer is not activated (Sanchez-Higueras et al., 2014). As VT03 expression is independent of Dfd and Scr and is sufficient to activate *snaR2P2* expression (Fig. 4G), the lack of *snaR2P2* activation in the glands of *Dfd Scr* mutant embryos must be due to a parallel Hox requirement and not caused by the lack of Upd1 expression. Because the *rep2-Upd1::HA* minigene can rescue tracheal formation, we next analysed if rep2 expression is affected in *Scr Antp Ubx* mutants where most functional Hox is absent in the trunk segments. In *Scr Antp Ubx rep2-GFP* embryos although the thoracic stripes are less defined (Fig. 5E-F) expression is maintained.

**Fig. 5.**
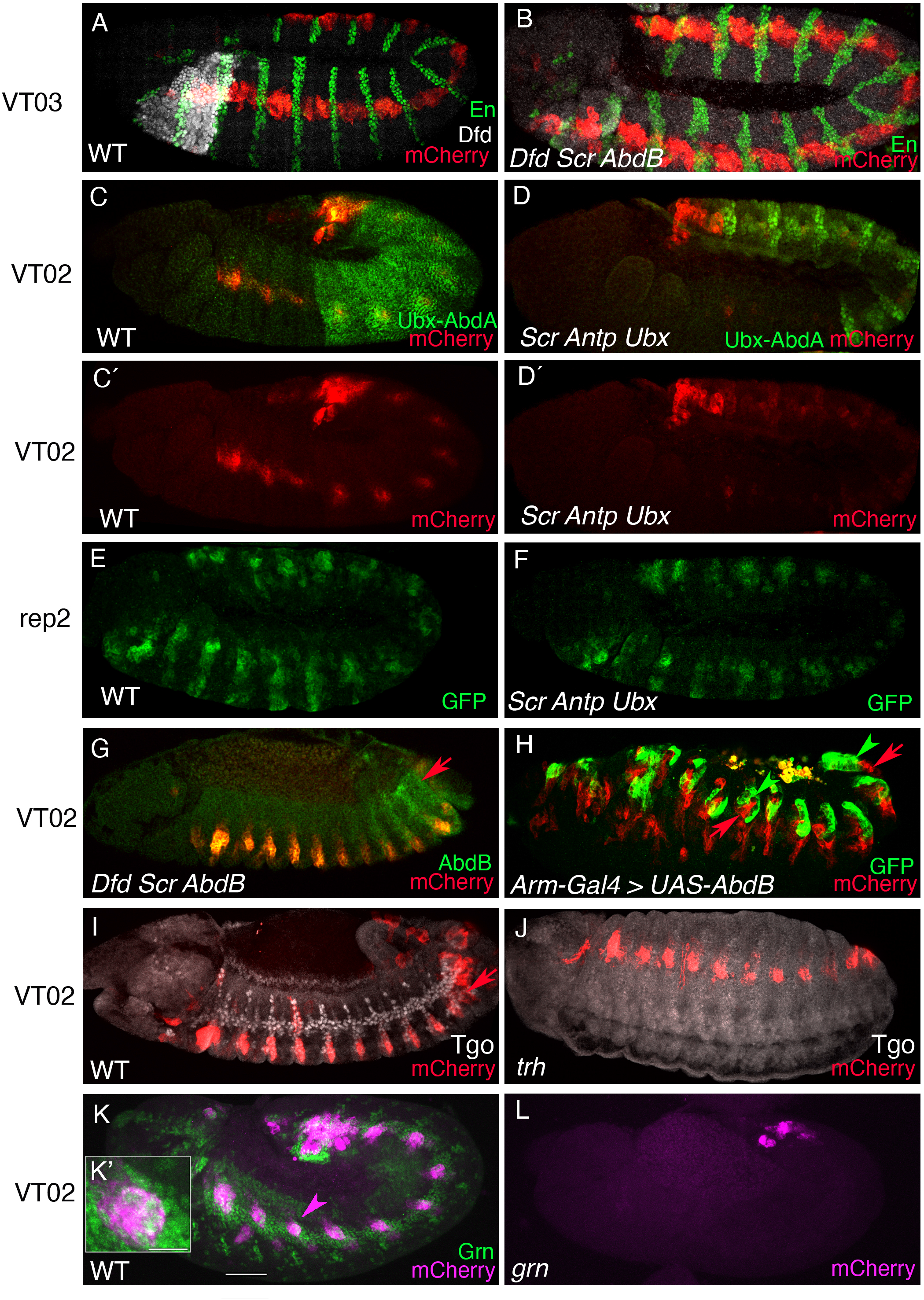
Regulation of *upd* organogenetic enhancers. (A-B) *VT03-mCherry* expression in a wild type (A) or a *Dfd Scr Abd-B* (B) embryo stained with anti-RFP (red), anti-En (green) and anti-Dfd (white) antibodies. (C-D) *VT02-mCherry* expression in a wild type (C, C’) or a *Scr Antp Ubx* (D, D’) embryo double stained with an antibody detecting Ubx and Abd-A. (E-F) *rep2-GFP* expression in a wild type (E) or a *Scr Antp Ubx* (F) embryo. (G) *VT02-mCherry* expression in a *Dfd Scr Abd-B* embryo double stained with anti-AbdB. This panel is a composite photo of two different planes to show the posterior spiracle site (red arrow) and the trachea primordia in the same image. Note the lack of VT02 expression in the posterior spiracle site. (H) *VT02-mCherry enD-GFP* embryos expressing ectopically *UAS-AbdBm* with *arm-Gal4* showing the ectopic expression of the *enD-GFP* stigmatophore enhancer associated to *VT02-mCherry* spiracular chamber expression. Red arrows point at the spiracular chamber and green arrowheads at the stigmatophore cells in A8 and in the A2 ectopic posterior spiracles. (I-J) *VT02-mCherry* expression in a wild type (I) or a *trh*^8^ (J) embryo double stained with anti-Tgo antibody to label the trachea. (K) *VT02-mCherry grn:GFP BAC* embryos show Grn is expressed in the *VT02-mCherry* cells. (K’) shows a close-up of the A1 segment (arrowhead). (L) *VT02-mCherry* expression only remains in the posterior spiracle of *grn^7J86^* mutant embryos. Scale bars 50 µm in K, and 10 µm in K’.

We also analysed the tracheal expression of *VT02-mCherry* in *Scr Antp Ubx* triple mutants where we observed reporter activation decreases in the trunk segments with the exception of A8 where Abd-B is still present (Fig. 5C-D). To test if the posterior spiracle expression is regulated by Abd-B, we studied *VT02* expression in *Dfd Scr Abd-B* embryos. We found that in homozygous mutants, A8 activation is absent from the posterior spiracle primordium although the tracheal associated expression remains (Fig. 5G). To confirm that *VT02-mCherry* spiracle regulation is downstream of Abd-B we combined it with *enD-GFP*, an *engrailed* posterior spiracle specific enhancer, that allows recognising *VT02-mCherry* spiracle expression from that associated to the trachea (Molina-Gil et al., 2023). In embryos where the *arm-Gal4* line ectopically activated *UAS-Abd-Bm* isoform all along the ectoderm, we detect ectopic *VT02* expression associated to the *enD* spiracle activity in all anterior segments (Fig. 5H). These results indicate that Hox input plays a secondary role on Upd regulation that is mostly confined to the VT04 and VT02 enhancers [(Barcenilla-Merino et al., 2025) and this work].

*VT02-mCherry* is expressed in the spiracular branches of the T3-A7 segments, which are the only main tracheal branches lacking a lumen in the larva and not contributing to gas exchange. However, these spiracular branches are functional, as they help to shed the tracheal cuticle during larval moulting and contribute to form the adult tracheal system after metamorphosis (Manning and Krasnow, 1993; Weaver and Krasnow, 2008). We find that VT02 tracheal expression is not affected in *trh* or in *trh vvl* double mutants, which are two of the main tracheal specification transcription factors (Fig. 5I-J), but disappears in *grn* mutants (Fig. 5K-L) that have been shown to be required for the formation of the proximal tracheal elements forming the spiracular branches connecting to the external epithelium (Matsuda et al., 2026; Matsuda et al., 2015).

## Discussion

During embryogenesis the JAK-STAT pathway is functionally required for segmentation and organogenesis (Hombria and Sotillos, 2013). With the aim of discovering when the specification of different organs by the JAK-STAT signalling pathway takes place, we have searched for enhancers regulating the transcription of its ligands and made tagged minigenes expressing a *upd1::HA* cDNA to test what defects caused by the deletion of all three *upd* ligands are rescued. We favoured this functional minigene approach over an enhancer deletion approach to solve the complication of the probable existence of redundant shadow enhancers in a locus that may have originated by tandem duplication. Flies carrying the minigene constructs were healthy and presented no detectable defects. By combining each of them with the *Dfos1A* deletion we have been able to pinpoint the functions rescued by each enhancer, discovering the precise temporal requirement of STAT for each embryonic organogenetic function (Table 1). Such minigene rescue approach has confirmed Upd1’s direct involvement in organogenesis, discarding that any of the observed organogenetic defects are caused indirectly by JAK-STAT’s earlier requirement to establish embryonic segmentation. Although our research has focused on the embryonic functions, the tools described here could be applied at later stages. As described above, *Df(1)os1A/ Y, Dp(1;3)DC344/+* and *Df(1)os1A/ Y; Dp(1;3)DC345/+* males have a normal embryogenesis, allowing future studies of larval or pupal functions analysing the BAC element rescues combined with or without minigenes.

### Location of organogenetic and gonadal enhancers

Guided by our analysis and previous genomic studies (Kvon et al., 2014) we have uncovered the location of the *upd* testis and ovary enhancers and most of the embryonic enhancers required for early organogenesis with the exception of the enhancer expressed in the embryonic eye primordium and the one driving expression in the clipeo-labrum (Figs. 1–2, summarized in Table 1). Of the enhancers analysed in this work none drive expression in the larval imaginal discs; and except *rep2,* that is expressed weakly in the blastoderm, and rescues the formation of segment A4 (Fig. 3E), none are involved in segmentation. The use of a minigene functional approach confirms that STAT is required during the specification and organogenesis of multiple organs including tentorium, endocrine glands, trachea, posterior spiracles and hindgut elongation independently of its earlier segmentation function.

### Parallel Upd and Hox interactions during organogenesis

We have previously shown that the specification of many of the organs mentioned above requires both JAK-STAT signalling and Hox gene input. At the molecular level this is shown by the simultaneous requirement of both Hox and JAK-STAT input for the activation through the *vvl1+2* enhancer of the *ventral veinless* gene in the metameric tentorium/glandular/tracheal primordia along the cephalic and trunk segments, the activation of the ring gland specific *sna* enhancer, and the posterior spiracle regulation of the *crumbs* gene (Barcenilla-Merino et al., 2025; Pinto et al., 2015; Sanchez-Higueras et al., 2019; Sanchez-Higueras et al., 2014). We have analysed the possibility that Hox gene function in these organs is mediated through the spatial regulation of early *upd* enhancers by testing VT and rep enhancer activity in various Hox mutant backgrounds. Our experiments show that Hox input is not required for *VT03* or *rep2* activation, confirming that for the formation of the trachea, and glands the STAT and Hox inputs act in parallel over common targets (Garcia-Ferres et al., 2022). A different situation arises with VT02 where trunk Hox genes are required for *upd1* expression in a subset of ventral cells in the tracheal pits and the posterior spiracles (Fig. 5D,G), and with VT04 in the head where Dfd and Scr are required for its activation in the maxillary and labial segments (Barcenilla-Merino et al., 2025). It would be interesting to find if Hox input on VT02 is mediated through *grn* regulation (Fig. 5L).

### Shade enhancers define a new class of redundant *cis*-regulatory elements

Shadow enhancers are quasi-redundant *cis*-regulatory elements activating gene expression in similar patterns and controlled by similar upstream factors. In laboratory conditions, redundant shadow enhancers have been shown to provide robustness to gene expression that may be especially important in different environmental conditions (Frankel et al., 2010; Perry et al., 2010), and have also been shown to contribute to expression pattern precision and evolvability [Reviewed in (Barolo, 2012; Kvon et al., 2021)]. As expected from its probable tandem duplication origin, the *upd* locus contains a series of shadow enhancers driving very similar expression, as illustrated by the many elements we have found to drive head cyst cell expression in the testis terminal region (Fig. 2A-F). Unexpectedly, we have found a new class of functionally redundant enhancers we name *shade enhancers* that despite having different expression patterns, are capable of performing the same functions and rescue the same organs. This is illustrated by *rep2-Upd::HA* and *VT03-Upd::HA* both of which can rescue tracheal specification despite one being expressed in segmental stripes of cells abutting the posterior compartment, and the other being expressed in a longitudinal antero-posterior stripe spanning most cephalic and trunk segments (Fig. 1D-E). Similarly, ring gland specification and migration can be rescued both by *VT03-Upd::HA* and by *VT04-Upd::HA* which are expressed in different patterns dependent on different upstream factors (Fig. 1C, E). We propose that many other genes may contain shade enhancers that have not been discovered due to the absence of systematic functional tests as the ones presented in this work. Shade, as shadow enhancers, could confer regulatory robustness and precision during development, but shade enhancers offer the unique characteristic of providing embryonic gene expression plasticity that could allow gene expression patterns to change during evolution without modifying the animal’s underlying body plan. Thus, unlike shadow enhancers, shade *cis*-regulatory elements could be important contributors to developmental system drift (McColgan and DiFrisco, 2024).

## Materials and methods

### Fly stocks

Flies were reared at 25°C on standard *Drosophila* medium. The following transgenic lines and mutant alleles from the Bloomington stock center were used:

*Reporter genes: vvl1+2-GFP*, *snaR2P2-GFP (Sanchez-Higueras et al., 2014)*, enD-GFP (Molina-Gil et al., 2023), *vvl-risk-lacZ (Sotillos et al., 2010)*, *btl-moeRFP (Caussinus et al., 2008)*.

*Mutant alleles: upd^YM55^* (B#4767)*; Df(1)os1A (Hombria et al., 2005)* ; *lab^14^* (B#2092); *Ubx^1^* (B#2866); *lab^CRISPRΔSev^ Dfd^6^ Scr^4^* (this work)*; Dfd^6^ Scr^4^ Abd-B^M1^*(recombined for this work); *Scr^C1^ Antp^NS+RC3^ Ubx^MX12^ (Struhl, 1982)*; *trh^8^* (Isaac and Andrew, 1996); *vvl^GA3^ trh^8^* (Sotillos et al., 2010); *grn^7J86^ (Brown and Castelli-Gair Hombria, 2000)*; *arm-Gal4*, *UAS-AbdBm (Rivas et al., 2013); grn::GFP BAC* (B#58483); *nos-Cas9* (B#54591).

*upd locus BAC elements:* Dp(1;3)DC343 (B#32277) coordinates 1:18070838..18152488; Dp(1;3)DC344/TM3Sb (stock not available in Bloomington any longer) coordinates 1:18241601..18324274; Dp(1;3)DC345 (B#30758) coordinates 1:18189748..18282159. All three BAC elements were recombined with *Ubx^1^* to label the chromosome carrying the BAC element.

### Upd BAC rescues

*Df(1)os1A/FM6 w ct* females were crossed to wild type, *Dp(1;3)DC343* or *Dp(1;3)DC345* homozygous males and cuticle preparations were analysed. No *upd* phenotypes were observed in the cuticle preparations of *Df(1)os1A/ FM6 w ct* females mated to *Dp(1;3)DC345* males although no adult *Df(1)os1A/ Y; Dp(1;3)DC345/+* males were obtained from the cross. Similar results were obtained after crossing *upd^YM55^/FM7* females to *Dp(1;3)DC345* males indicating this BAC duplication rescues the embryonic phenotypes but not all larval or pupal functions resulting in lethality. To make sure that the embryo cuticles we are analysing carry the different BAC duplications, we repeated the crosses with all three duplications recombined with *Ubx^1^*. For that purpose we crossed *Df(1)os1A/ FM6 w ct* females to *Ubx^1^/TM6B* males and the resulting *Df(1)os1A/+; Ubx^1^/+* females were mated to *Dp(1;3)DC343 Ubx^1^/TM6B*, *Dp(1;3)DC344 Ubx^1^/TM6B* or *Dp(1;3)DC345 Ubx^1^/TM6B*males. The resulting *Ubx* homozygous embryos (recognizable because the first abdominal denticle belt acquires thoracic morphology) only showed *upd* defects in the *Dp(1;3)DC343 Ubx^1^/TM6B* crosses, confirming that *Dp(1;3)DC344* and *Dp(1;3)DC345* can rescue the embryonic phenotypes.

### Generation of a CRISPR *labial* mutation on a *Dfd Scr* stock

CRISPR/Cas9 mutation of *labial* was performed using gRNAs cloned into the *pCFD5* vector, following the published pCFD5 cloning protocol for multiplex gRNA expression (Port et al., 2014). The gRNA-containing oligos were designed according to the pCFD5 tRNA-flanked gRNA strategy and cloned into the vector as described in that protocol. The *labial* guide sequences used were CAGCTGGACATGTGCCGCGG and GAATGAGGCCCTCCTTCACG.

The pCFD5-gRNA plasmid DNA was sent to CBM-SO Madrid for embryo injection and generation of transgenic flies in a *white* genetic background. The resulting transgenic lines were crossed to a *nos-Cas9* stock (Port et al., 2014) and subsequently combined with the *Dfd^16^ Scr^4^* mutant background. Males displaying the characteristic small sex combs phenotype of *Scr* were selected and crossed to *lab^14^/TM3* mutant flies. Crosses where no *lab* homozygous adult flies appeared were kept as stock. Confirmation of a precise deletion of the targeted *labial* homeobox region was confirmed by PCR using the forward 5′-TTCCTTGATTGATCTTACCCCTTAGTGC-3′ and reverse 5′- TTGTATGTATTTATTTCCTCTCTGCACTTTCTACTA-3′ primers followed by Sanger sequencing.

### Cuticle preparations

Embryos were collected overnight on agar apple juice plates supplemented with yeast and allowed to age for 24h. Embryos that remained in the chorion were collected by hand, dechorionated in 50% commercial bleach, washed and transferred to a microscope slide. After absorbing the excess of water with a tissue paper, embryos were transferred into a drop of Hoyer’s medium. After spreading the embryos, they were covered with a coverslip and subjected to mild pressure to crack the vitelline membrane. The slides were incubated on a 75°C hot plate for 48 hours with a small weight on top. Cuticles were analysed and photographed in an Axioplan2 Zeiss microscope using phase contrast.

### Reporter constructs

*VT02-mCherry, VT03-mCherry, VT04-mCherry*: Using the information available at the Stark lab (https://enhancers.starklab.org) we selected three intergenic regions driving an overlapping expression with *upd1*. *VT63702* (here named *VT02* for simplification) is active in the trachea, *VT63703* (*VT03*) in an antero-posterior stripe of cells, and *VT63704* (*VT04*) in the cephalic region and hindgut. Using specific primers adding a BamHI restriction site at the end (Table 2) we amplified by PCR these elements from genomic DNA. The amplified elements were sequenced and cloned in the BamHI site of pCaSpeR-mCherry-PH previously made in the laboratory.

**Table 2:**
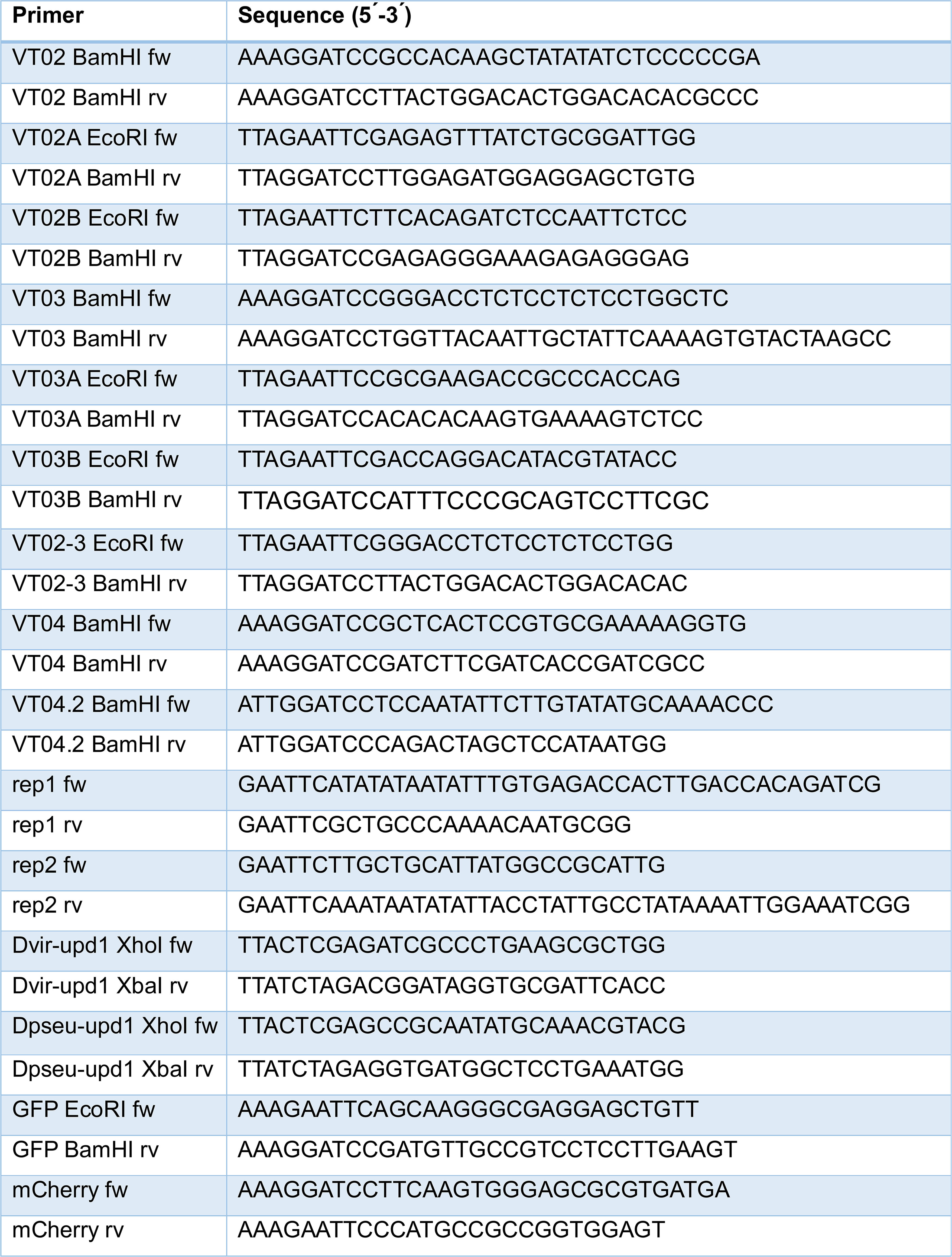
Oligos used for reporter constructs and RNA *in situ* probes.

*VT02A-GFP, VT02B-GFP, VT03A-GFP, VT03B-GFP, VT02-3-GFP*: The VT02 and VT03 enhancers were subdivided giving rise to VT02A and VT02B, and VT03A and VT03B taking care not to break sequence blocks conserved in several *Drosophila* species as detected using the USCS Genome Browser, http://genome.ucsc.edu/ (Sup. Fig. 2). VT02-3 was made from the overlapping VT02 and VT03 region. The 5 fragments were obtained by PCR using the plasmids pCasper mCherry PH VT63702 and pCasper mCherry PH VT63703 as templates (the first one for constructs VT02A, VT02B and VT02-3, and the second one for constructs VT03A and VT03B). The fragments range from 449 to 740bp in length. PCR reactions with specific oligos (Table 2) were performed with GoTaq G2 Flexi DNA polymerase (Promega) and fragments were cloned into pGEM-T Easy vector. After sequencing confirmation, fragments were digested with EcoRI and BamHI enzymes (sites added in the 5’ ends of the primers) and subcloned into pCasper eGFP-PH vector digested with the same restriction enzymes, except for VT03B, which was digested with BamHI and SpeI enzymes.

*VT04.2-GFP*: We amplified VT04.2 by PCR from the pCasper-mCherry-PH-VT04 plasmid using the GoTaq DNA polymerase and specific primer pairs (Table 2). The PCR product was subcloned in a pGem-T Easy vector. After sequencing confirmation, we cloned the BamHI fragments into the pCaSper-mCherry-PH reporter plasmid. *rep1-GFP, rep2-GFP*: For *rep1* and *rep2* enhancer reporters, the enhancer fragments were amplified using specific primers (Table 2) and cloned into the *pCasperEGFP* reporter vector (Sanchez-Higueras et al., 2014). These constructs were used exclusively for enhancer reporter assays and not for minigene generation.

**Minigenes** *VT02-Upd1::HA*, *VT03- Upd1::HA*, *VT04-Upd1::HA*, *rep2-Upd1::HA*.

Minigene constructs containing *upd1::3xHA* were generated by subcloning the VT02, VT03 and VT04 enhancers into the *pCasper-hs43-polylinker* vector using the *BamHI* restriction site (Sanchez-Higueras et al., 2014). The *upd1::3xHA* fragment was synthesized by BIOMATIK and cloned into the *KpnI* site of the resulting vector in the correct orientation. For the rep2 minigene construct, the *upd1* cassette was first cloned into the *KpnI* site of the *pCasper-hs43-polylinker* vector. Subsequently, the rep2 enhancer was cloned into the *EcoRI* site of the vector. The rep2 enhancer fragment was PCR-amplified with primers described in Table 2.

All constructs were injected by the CBM *Drosophila* transgenesis service (Spain). Between 2 and 6 independent random inserts were analysed for each construct.

### RNA *in situ* hybridization

*upd1* probes were prepared by PCR amplification using genomic DNA from *Drosophila virilis* or *D. pseudoobscura* as templates. Primers were designed to amplify the *D. virilis* and *D. pseudoobscura* sequences orthologous to the *D. melanogaster upd1* gene (Table 2).

We used GoTaq G2 Flexi DNA polymerase (Promega). PCR products were digested with XhoI and XbaI restriction enzymes (sites added in the 5’ ends of the primers) and ligated into pBluescript II SK vector (Stratagene) digested with the same enzymes. After sequencing confirmation, constructs were digested with XhoI, and transcribed *in vitro* using the DIG RNA Labelling Kit (Roche) with T3 polymerase from the T3 promoter to obtain the antisense probes. The probes’ sizes are 195bp for *D. virilis* and 322bp for *D. pseudoobscura*.

The *mCherry* probe was obtained from a pCaSpR-mCherry plasmid. Adapter sequences for T7 and T3 transcription enzymes were added to the primers for amplification via PCR. The mCherry insert was then cloned into a pBluescript plasmid (Stratagene), sequenced and *in vitro* transcribed using the DIG RNA Labelling Kit (Roche) to generate the anti-sense strand for the mCherry sequence. The *GFP* probe was made by PCR using specific oligos (Table 2). NBT-BCIP was used to stain the sheep anti-DIG-AP.

### Immunocytochemistry

Embryos were dechorionated in 50% commercial bleach for 5 minutes, fixed in a mixture of heptane: 4% formaldehyde in PBS for 20 minutes. After removing the formaldehyde lower phase, methanol was added and the embryos shaken vigorously to break the vitelline membrane. Sinking embryos were recovered and washed in fresh methanol. Fixed embryos were rehydrated in PBS with Tween 0,1% and washed tree times in PBS with Tween 0,1% and BSA 1% followed by overnight incubation at 4^◦^C with primary antibodies diluted in PBS with Tween 0,1% BSA 1%. Embryos were washed three times in PBS with Tween 0,1% BSA 1% and incubated 2 hours at room temperature (RT) in PBS with Tween 0,1% BSA 1% with the secondary antibodies. Finally, embryos were rinsed three times in PBS with Tween 0,1% and washed twice with PBS before mounting in Vectashield (H-1000, lot.nr: ZL0603).

For testes and ovaries five-day old adults were dissected in PBS and fixed for 30 minutes at RT in 4% paraformaldehyde in PBS+0.3% Triton X-100, washed three times with PBS-Triton 0.3% and preadsorbed in PBS-Triton 0.3%+ 1%BSA during 1 hour. Primary antibodies were incubated overnight at 4°C, washed four times in PBS-Triton 0.3%. Gonads were preabsorbed in PBS-Triton 0.3%+ 1%BSA for 1hour at RT. Secondary antibodies diluted at 1:400 in PBS-Triton 0.3%+ 1%BSA were incubated at RT during 4 hours. After incubation, gonads were washed 4 times 15 minutes in PBS-Triton 0.3% at RT, rinsed twice in PBS and mounted in Vetacshield.

Antibodies and markers used are described in Table 3.

**Table 3:**
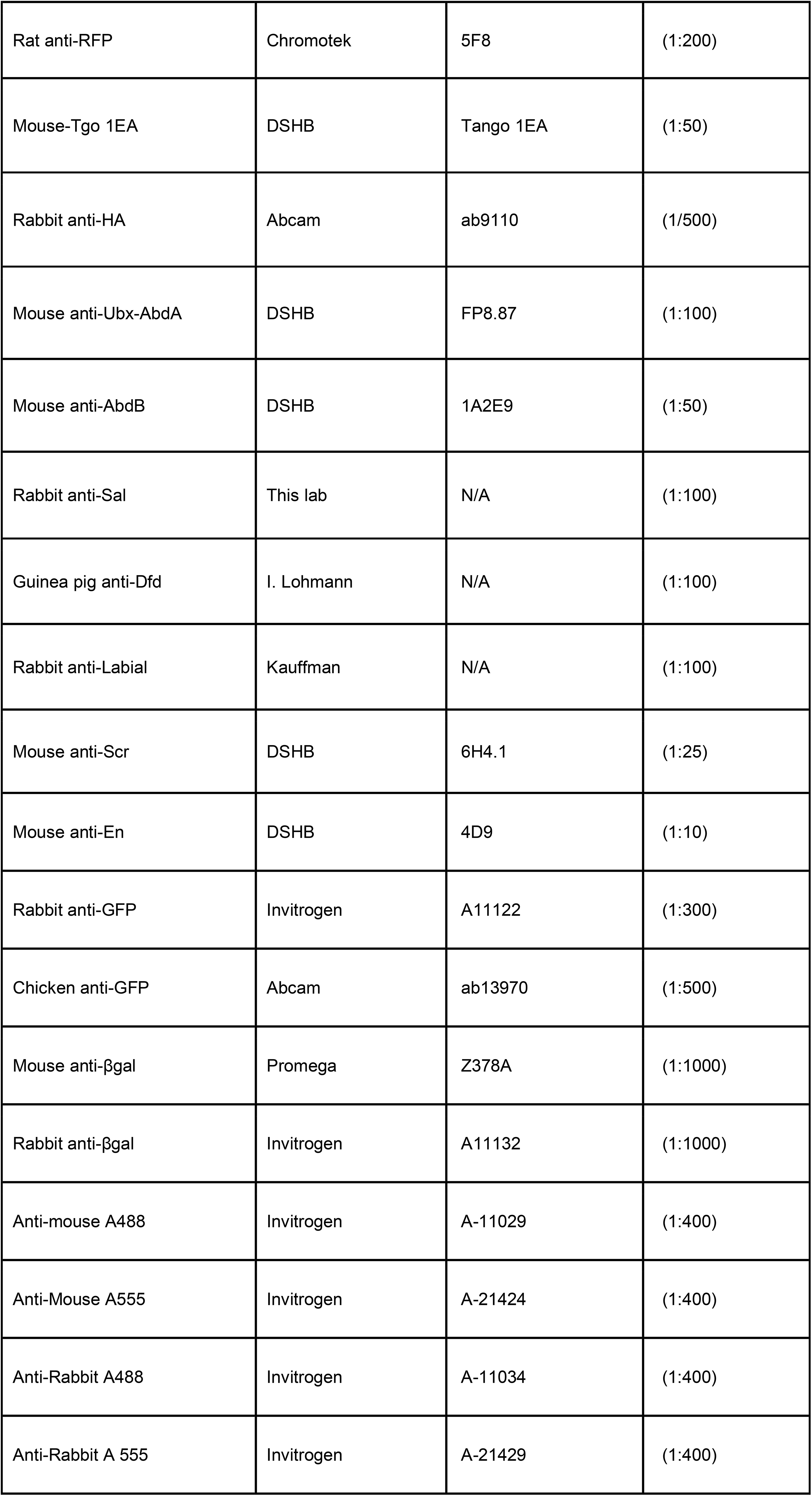
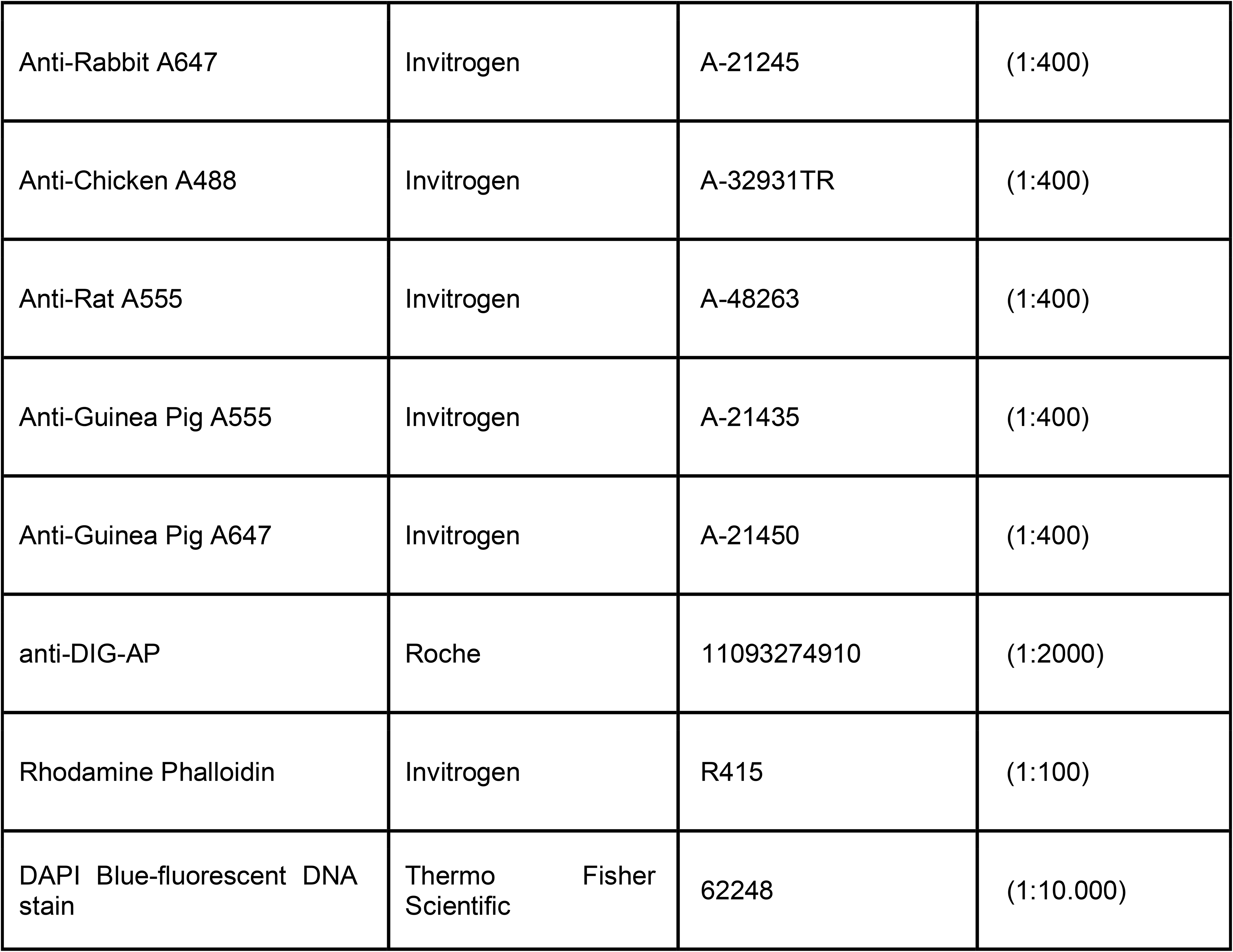
Antibodies and markers.

### Imaging

Confocal images were taken in an SPE or a Stellaris Leica microscope. Images were treated with ImageJ and Photoshop. RNA *in situ* and cuticle preparations were photographed with an Axioplan2 Zeiss microscope.

## Acknowledgements

We thank Jose María Urbano for critical reading of the manuscript. This work was supported by Ministerio de Ciencia e Innovación grants PID2019-104656GB-I00, PID2022-136728NB-I00 and PID2025-171772NB-I00 co-funded by the European Regional Development Fund (FEDER) to J.C.-G.H and S.S and a grant BES2017-081120 to MGF. The CABD was supported by a María de Maeztu Unit excellence grant CEX-2020-001088-M.

**Sup. Fig. 1. *upd* RNA and enhancer activation in dorsal neuroblasts and tracheal branch subsets.**

(A) Close up of a st9 embryo showing *upd* RNA is expressed along a continuous row of neuroblasts as well as in segmental stripes. (B) *VT03-mCherry* st11 embryo carrying the *vvl1+2-GFP* tracheal marker shows persistent mCherry expression overlaps a subset of ventral cells in the tracheal primordia (arrow). (C) Ventral view of a *VT03-mCherry* embryo showing persistent mCherry expression labelling the ventral tracheal (vt) branches and some axons crossing the ventral midline (in). (D) Lateral view of a st13 *VT03-mCherry lab Dfd Scr* mutant showing cephalic expression unaffected. (E) Close up of a *VT02-mCherry vvl1+2-GFP* st11 embryo showing expression in the posterior spiracle (ps). (F) *VT02-mCherry vvl1+2-GFP* stage 12 embryo showing expression in the spiracular branches (sb) and posterior spiracles. (G) *VT02-mCherry* embryo stained with the tracheal marker Tgo (green) shows expression in the tentorium (t), anterior spiracle (as), posterior spiracle (ps) and tracheal spiracular branches (sb). (H) Ventro lateral view of a st13 *VT03-mCherry lab Dfd Scr* mutant showing persistent expression in ventral trachea (vt) and axons crossing the midline (in). Scale bar 50 µm.

**Sup. Fig. 2. Schematic location of the *cis*-regulatory fragments analysed**

Position of the different *cis*-regulatory fragments analysed and sequence conservation of the *upd1* locus obtained using the USCS Genome Browser, http://genome.ucsc.edu/. For simplification VT63702, VT63703 and VT63704 are referred throughout the text as VT02, VT03 and VT04. The approximate location of each enhancer is shown underneath. The position of the HCC enhancers is shown relative to the large 2kb fragments, while other enhancers are mapped with respect to the sub-fragments.

**Sup. Fig. 3. Drosophila virilis and D. pseudobscura upd1 RNA expression.**

Left column (A) shows *D. virilis upd1* RNA expression, and right column (B) shows *D. pseudobscura upd* RNA expression. At equivalent stages RNA expression is indistinguishable from *D. melanogaster upd1* transcription.

**Sup. Fig. 4. *Upd1::HA* minigene expression.**

(A-Á’) *VT02-Upd1::HA* minigene expression. St14 embryo stained with anti-HA (A), and double stained (Á) with anti-HA (green) and anti-En (purple) showing expression in the anterior spiracle (as) and in the spiracular branches (sb). (Á’) Close up of embryo in A’. (B) *VT04-Upd1::HA* minigene. St12 embryo stained with anti-HA (green) showing expression in the tentorium/gland primordia. Scale bar in B 50 µm.

**Sup. Fig. 5. Expression of *upd* enhancers in *Dfos1A* mutants lacking all *upd* ligands.**

(A) Control *rep2-GFP* embryo stained with anti-GFP (green) and anti-En (purple) showing expression in an adjacent stripe posterior to the En labelled cells. (B-E) *Dfos1A* mutant embryos carrying *rep2-GFP* (B), *VT02-mCherry* (C), *VT03-mCherry* (D), or *VT04-mCherry* (E). Embryos in B-D are double stained with anti-En to show the embryonic segment defects. (D’) presents a z-section showing the reporter and the engrailed cells are in the same plane, indicating that the enhancer expressing cells normally forming tracheal branches have not invaginated in *Dfos1A* embryos. (A-B) st11 embryos, (C-E) st14 embryos. Scale bar 50 µm.

